# Exogenous mitochondrial transfer increases energy expenditure and attenuates adiposity gains in mice with diet-induced obesity

**DOI:** 10.1101/2023.12.23.573206

**Authors:** Maria Namwanje, Soumi Mazumdar, Amanda Stayton, Prisha S. Patel, Christine Watkins, Catrina White, Chester Brown, James D. Eason, Khyobeni Mozhui, Cem Kuscu, Navjot Pabla, Erin J. Stephenson, Amandeep Bajwa

**Author notes:** Co-corresponding authors Addresses for correspondence: Amandeep Bajwa, Ph.D, Associate Professor, Transplant Research Institute, Department of Surgery, University of Tennessee Health Science Center, 71 S Manassas St, Room 418H, Memphis, TN 38103, Erin Stephenson, Ph.D., Assistant Professor, Department of Anatomy, College of Graduate Studies, Midwestern University, 555 31^st^ Street, Suite 542-L, Downers Grove, IL 60515.

## Abstract

Obesity is associated with chronic multi-system bioenergetic stress that may be improved by increasing the number of healthy mitochondria available across organ systems. However, treatments capable of increasing mitochondrial content are generally limited to endurance exercise training paradigms, which are not always sustainable long-term, let alone feasible for many patients with obesity. Recent studies have shown that local transfer of exogenous mitochondria from healthy donor tissues can improve bioenergetic outcomes and alleviate the effects of tissue injury in recipients with organ specific disease. Thus, the aim of this project was to determine the feasibility of systemic mitochondrial transfer for improving energy balance regulation in the setting of diet-induced obesity. We found that transplantation of mitochondria from lean mice into mice with diet-induced obesity attenuated adiposity gains by increasing energy expenditure and promoting the mobilization and oxidation of lipids. Additionally, mice that received exogenous mitochondria demonstrated improved glucose uptake, greater insulin responsiveness, and complete reversal of hepatic steatosis. These changes were, in part, driven by adaptations occurring in white adipose tissue. Together, these findings are proof-of-principle that mitochondrial transplantation is an effective therapeutic strategy for limiting the deleterious metabolic effects of diet-induced obesity in mice.

## Introduction

Transfer of mitochondria between cell types is increasingly recognized as being important in the regulation of tissue structure and function^1-6^. In conditions associated with mitochondrial dysfunction, local or systemic transfer of exogenous mitochondria from healthy donor tissue has been shown to improve bioenergetic outcomes and alleviate the effects of tissue injury in recipients^7-14^. Given obesity is associated with chronic multi-system bioenergetic stress, the aim of this project was to determine the feasibility of systemic mitochondrial transfer for improving energy balance regulation in the setting of diet-induced obesity. We found that transplantation of mitochondria from lean mice into mice with diet-induced obesity attenuated adiposity gains by increasing energy expenditure and promoting the mobilization and oxidation of lipids. Additionally, mice that received exogenous mitochondria demonstrated improved glucose uptake, greater insulin responsiveness, and complete reversal of hepatic steatosis. These changes were, in part, driven by adaptations occurring in white adipose tissue. Together, these findings are proof-of-principle that mitochondrial transplantation is an effective therapeutic strategy for limiting the deleterious metabolic effects of diet-induced obesity in mice.

## Results

After validating that exogenous mitochondria are taken up into NIH-3T3-L1 cells in a dose-dependent manner *in vitro* (Extended Data 1a), we observed that cells treated with exogenous mitochondria have increased respiration (Extended Data 1b), demonstrating that ADP-dependent respiration increases with increasing mitochondrial dose (Extended Data 1c). To determine whether addition of exogenous mitochondria could modify lipid metabolism in vitro, we treated NIH-3T3-L1 adipocytes with either vehicle (PBS) or 10 μg/well of exogenous mitochondria for up to 72 h (Extended Data 1d). Adipocytes treated with exogenous mitochondria had reduced lipid accumulation (Extended Data 1e) which was associated with increased expression of transcripts that encode proteins involved in energy dissipation and decreased expression of transcripts that encode proteins involved in lipogenesis and adipogenesis (Extended Data 1f, Extended Data Table 1). We also observed elevated basal lipolysis, and potentiated lipolysis following treatment with the β_3_-adrenegic agonist CL316243 in adipocytes treated with exogenous mitochondria compared to vehicle-treated cells (Extended Data 1g-h). Together, these findings suggest that transfer of exogenous mitochondria into NIH-3T3-L1 adipocytes *in vitro* increases the mobilization and oxidization of intracellular lipids, resulting in reduced lipid accumulation and, subsequently, smaller adipocytes.

To confirm that systemic administration of exogenous mitochondria results in broad tissue uptake of donor mitochondria, we injected 20 μg/g body weight of mitochondria (or vehicle) into the retro-orbital sinus of lean adult C57BL/6J male mice and collected and measured the epifluorescence of a selection of organs and tissues after 30 min, 2 h, 24 h, 48 h, or 72 h (Extended Data 2A). Uptake of exogenous mitochondria was observed in most tissues, including adipose tissue depots, as early as 30 min after injection and persisted up to 48 h (Figure 1a; Extended Data 2b-i). These data indicate that systemic administration of exogenous mitochondria is a feasible strategy for increasing the mitochondrial content of multiple tissues simultaneously *in vivo*.

**Figure 1:**
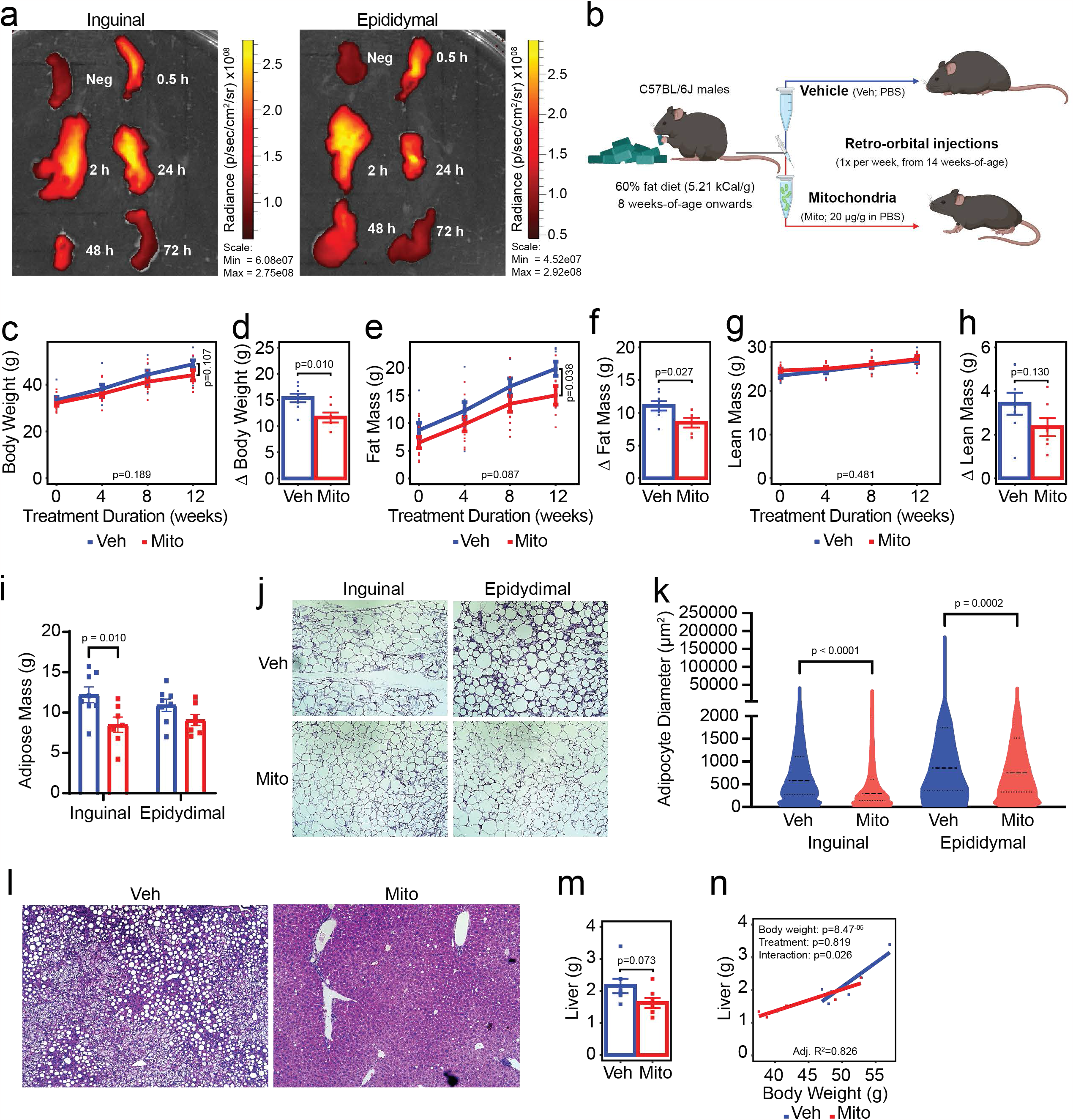
Transfer of exogenous mitochondria into mice with preestablished obesity attenuates subsequent high fat diet-induced adiposity gains. Fluorescent donor mitochondria are taken up into inguinal and epididymal adipose tissue depots (a). Schematic of experimental design (b). Effect of mitochondrial transfer (Mito; red lines/bars/points) compared to vehicle treatment (Veh; blue lines/bars/points) on body weight (c-d), fat mass (e-f), lean mass (g-h), individual adipose depot masses (i), adipocyte size (j-k) and liver steatosis (l) and weight (m-n). Values shown are the mean ± standard error with biologically independent replicates overlayed (n=7-8 Veh and n=8-10 Mito). Data were compared by linear mixed-effects models with each individual mouse fitted as a random intercept and treatment as the main fixed variable of interest (line graphs), Student’s t-test (bar graphs, 12-week time points on line graphs, and violin plots), or analysis of covariance, with body weight and treatment as covariates (scatter plot with regression lines). Refer to each figure panel for individual p-values. Note that the adjusted R^2^ value depicted in panel n reflects the proportion of variance resulting from treatment that can be explained by differences in body weight corrected for the number of variables included in the model (adjusted R^2^; multiple R^2^ = 0.866). Source data are provided in the source data file.

To determine if weekly mitochondrial transfers could mitigate the effects of diet-induced obesity on body composition, energy balance, and other related metabolic parameters we performed the following study: Beginning at eight weeks of age, male C57BL/6J mice were fed a high-fat diet (HFD; 60% kCal from fat, 5.21 kCal/g) for six weeks to induce obesity. At 14 weeks of age, mice began receiving once weekly intravenous injections of either exogenous mitochondria (Mito; 20 μg/g body weight) or vehicle (Veh; volume-matched PBS) while being maintained on HFD for an additional 15-week period (Figure 1b). After 12 weeks of treatment, mice receiving weekly mitochondrial transfers had gained less body weight than mice receiving vehicle (Figure 1c-d; -24.3%, p=0.010). The effect of mitochondrial transfer on body weight was due to attenuated adiposity gains (Figure 1e-f; -23.2%, p=0.027), without significant changes in lean mass (Figure 1g-h; p=0.130). Reduced adiposity in mice receiving mitochondrial transfers was consistent with these mice having less inguinal white adipose tissue than mice that received vehicle treatment (Figure 1i; -30.39%, p=0.010). Adipocytes in both the inguinal and epididymal white adipose depots were also smaller following treatment with mitochondria (Figure 1j-k; p<0.0001 and p=0.0002, respectively). Consistent with a reduction in whole-body adiposity, evidence of hepatic steatosis was completely absent in mice that received mitochondria compared to the vehicle group (Figure 1l; representative images), and there was a trend toward reduced overall liver weight (Figure 1m; -24.6%, p=0.073). When liver size was considered alongside body weight, 82.6% of the variance could be explained by the difference in body weight between the groups (Figure 1n; p=8.47^-05^) with treatment alone having no effect (p=0.819), despite there being a body weight-treatment interaction (p=0.026).

To determine which component of energy balance was driving the reduction in adiposity gains observed in mice receiving exogenous mitochondria, we housed mice in metabolic chambers that allow for simultaneous measurement of respiratory gases, food intake, and physical activity after 13-weeks of treatment with mitochondria (or vehicle). Compared to the mice receiving vehicle, mice that received mitochondrial transfers had elevated energy expenditure across both the light and dark photoperiods (Figure 2a; p=3.973^-05^ and p=2.468^-04^, respectively). Higher energy expenditure in mice receiving mitochondria was driven by increases in both resting and non-resting energy expenditure (Figure 2b; 20.2%, p=5.840^-04^ and 42.4%, p=0.012, respectively), which together contributed an additional 3.2 kCal/d of energy expenditure compared to vehicle treated mice (Figure 2c; p=3.820^-04^). Elevated non-resting energy expenditure was due, in part, to mitochondria-treated mice being more ambulatory during the dark photoperiod (Figure 2d; p=3.884^-05^), with increased ambulatory activity driving an increase in total activity compared to vehicle-treated mice (Figure 2e; 90%, p=6.00^-03^). No differences were observed between groups for absolute food intake over either photoperiod (Figure 2f), nor were any differences observed in total daily energy intake (Figure 2g) indicating the effect of mitochondrial transfer on adiposity was driven primarily by the increases in energy expenditure associated with both resting metabolism and additional energy expended as a result of increased physical activity.

**Figure 2:**
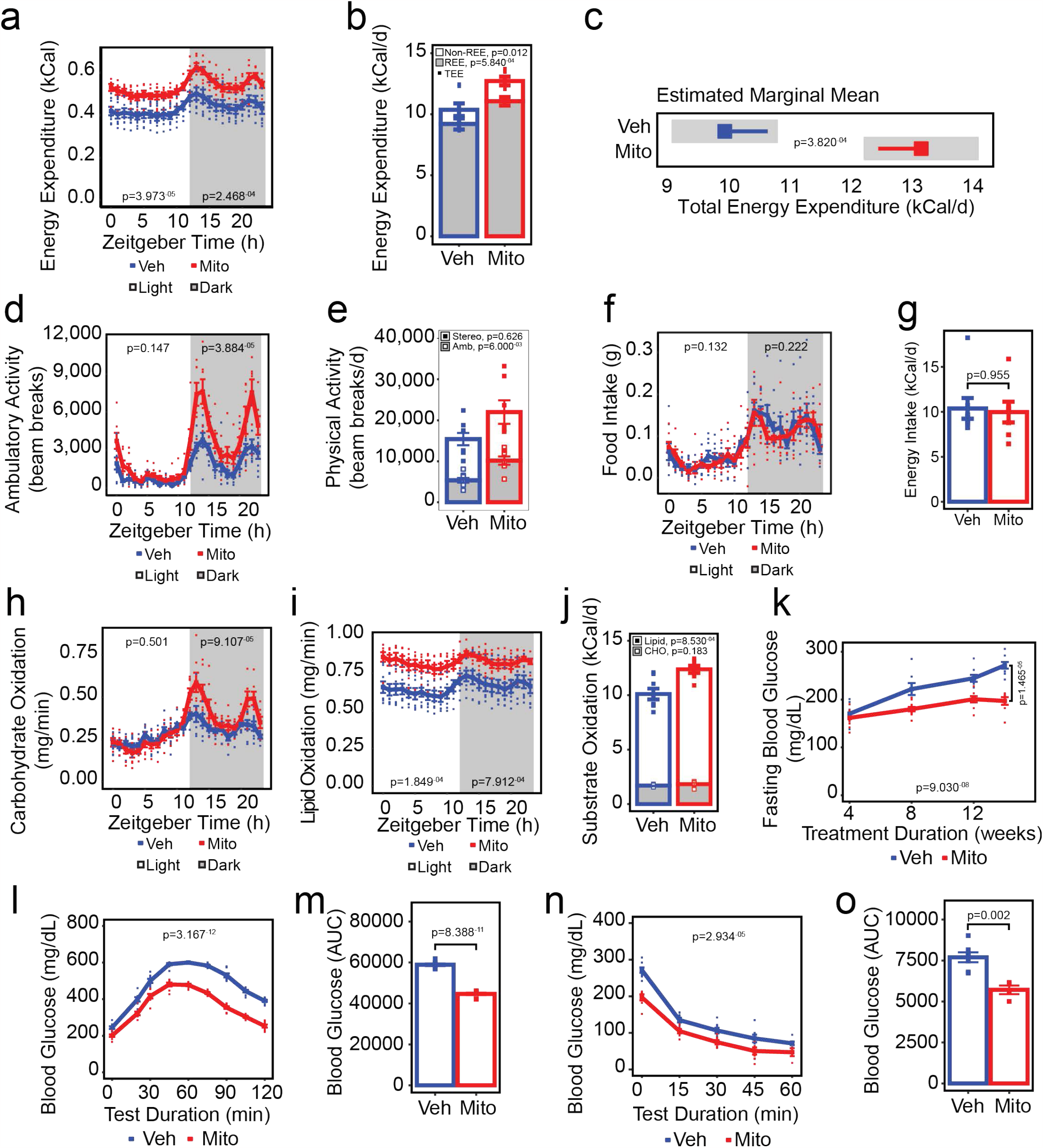
Transfer of exogenous mitochondria increases energy expenditure and enhances lipid oxidation and glucose tolerance in vivo. Effect of mitochondrial transfer (Mito; red lines/bars/points) compared to vehicle treatment (Veh; blue lines/bars/points) on energy expenditure (a-c), physical activity (d-e), food and energy intake (f-g), carbohydrate and lipid oxidation (h-j), fasting blood glucose concentrations (k), glucose tolerance (l-m), and insulin responsiveness (n-o). Values shown are the mean ± standard error with biologically independent replicates overlayed (n=8 Veh and n=7-10 Mito). In panel c, the grey bars represent the 95% confidence intervals. Data were compared by linear mixed-effects models with each individual mouse fitted as a random intercept and treatment as the main fixed variable of interest with (a, d, f, h, i) or without (k, l, n) fat mass and lean mass included in the models, Student’s t-test (bar graphs, 12-week time points on k), or analysis of covariance with lean mass and fat mass as covariates (estimated marginal means plot, c). Refer to each figure panel for individual p-values. Source data are provided in the source data file. Abbreviations: REE, resting energy expenditure; TEE, total energy expenditure; Stereo, stereotypic activity; Amb, ambulatory activity; CHO, carbohydrate.

To establish whether the energy expenditure-driven attenuation of adiposity gains was associated with specific changes in macronutrient oxidation, we leveraged the indirect calorimetry data to determine rates of carbohydrate and lipid oxidation (Figure 2h-j). In line with our observation that mitochondria-treated mice had increased dark photoperiod activity, we observed an increase in the rate of dark photoperiod carbohydrate oxidation (Figure 2h; p=9.107^-05^). Additionally, lipid oxidation rates were increased in mice receiving exogenous mitochondria compared to vehicle-treated mice during both the light and dark photoperiods (Figure 2i, p=1.849^-04^ and 7.912^-04^, respectively), suggesting that attenuation of adiposity gains in mitochondria-treated mice was due, in part, to increased oxidation of lipid substrates. Indeed, the proportion of energy expended as a result of the oxidization of lipid substrates was 36.56% greater in mice treated with exogenous mitochondria compared to those treated with vehicle (Figure 2j; open bars, p=8.530^-04^), whereas despite dark photoperiod increases in the rate of carbohydrate oxidation in mitochondria-treated mice, the relative contribution of carbohydrate oxidation to daily energy expenditure was not different between groups (Figure 2j, shaded bars; p=0.183).

To determine whether mitochondrial transfer was associated with changes in glucose homeostasis, fasting blood glucose concentrations were measured after four weeks of treatment onward (Figure 2k), whereas glucose and insulin tolerance tests were performed after 12- and 14-weeks of treatment, respectively (Figure 2l-o). Treatment with exogenous mitochondria lowered fasting glucose concentrations (Figure 2k; p=9.030^-08^), with mice that received exogenous mitochondria having 27.44% lower fasting glucose concentrations than mice that received vehicle after 14-weeks (p=1.465^-05^). Glucose tolerance was also improved in mice that received exogenous mitochondria (Figure 2l; p=3.167^-12^) with a 24.13% reduction in the area under the glucose curve during the glucose tolerance test compared to vehicle treated mice (Figure 2m, p=8.388^-11^). Notably, when differences in fasting blood glucose were considered, the effect of mitochondrial treatment on glucose tolerance persisted (p=1.239^-09^). Insulin responsiveness also improved in mice that received mitochondrial transfers (Figure 2n; p=2.934^-05^), with a 25.82% reduction in the area under the glucose curve compared to vehicle-treated mice during the insulin tolerance test (Figure 2o; p=0.002). However, improvements in insulin responsiveness were likely caused by differences in fasting glucose concentrations at the beginning of the test, as the groups were no longer different when fasting glucose was included as a fixed effect in the model (p=0.188).

Given that enhanced lipid metabolism appears to be a key driver of many of the metabolic adaptations we observed following exogenous mitochondrial transfer and our initial *in vitro* findings that demonstrate transfer of exogenous mitochondria increased respiration and lipolysis and reduced the lipid content of adipocytes *in vitro*, we sought to identify how mitochondrial transfer might modify adipose tissue directly. Similar to our *in vitro* findings, administration of exogenous mitochondria to mice with preexisting obesity led to changes in the expression of multiple transcripts across white adipose tissue depots. Transcripts that encode proteins involved in energy dissipating processes, lipolysis, fatty acid oxidation and autophagy were increased in both inguinal and epididymal depots from mice treated with exogenous mitochondria compared to vehicle-treated mice (Figure 3a-b, Extended Data Table 1). Transcriptional data indicating that mitochondrial treatment increases adipose tissue lipolysis were supported by the observation that compared to vehicle treated mice, mice that received exogenous mitochondria had increased serum non-esterified fatty acid and free glycerol concentrations (Figure 3c-d; 82.7%, p=0.0002 and 89.1%, p=0.0002, respectively), an adaptation likely required to support the increase in systemic lipid oxidation we observed in these mice. Conversely, several transcripts that encode proteins involved in *de novo* lipogenesis, fatty acid re-esterification, and inflammation were decreased in both inguinal and epididymal depots in mice that received mitochondria compared to vehicle-treated mice (Figure 3a-b, Extended Data Table 1), suggesting that white adipose tissue adaptation to increased mitochondrial content is not limited to changes that facilitate an increase in systemic fatty acid availability and oxidation. These data demonstrate that transfer of exogenous mitochondria into mice with preexisting obesity reprograms white adipose tissue to support elevated systemic lipid oxidation and increase overall energy expenditure while reducing the lipid storage and inflammatory signaling typically associated with high fat diet-induced obesity.

**Figure 3:**
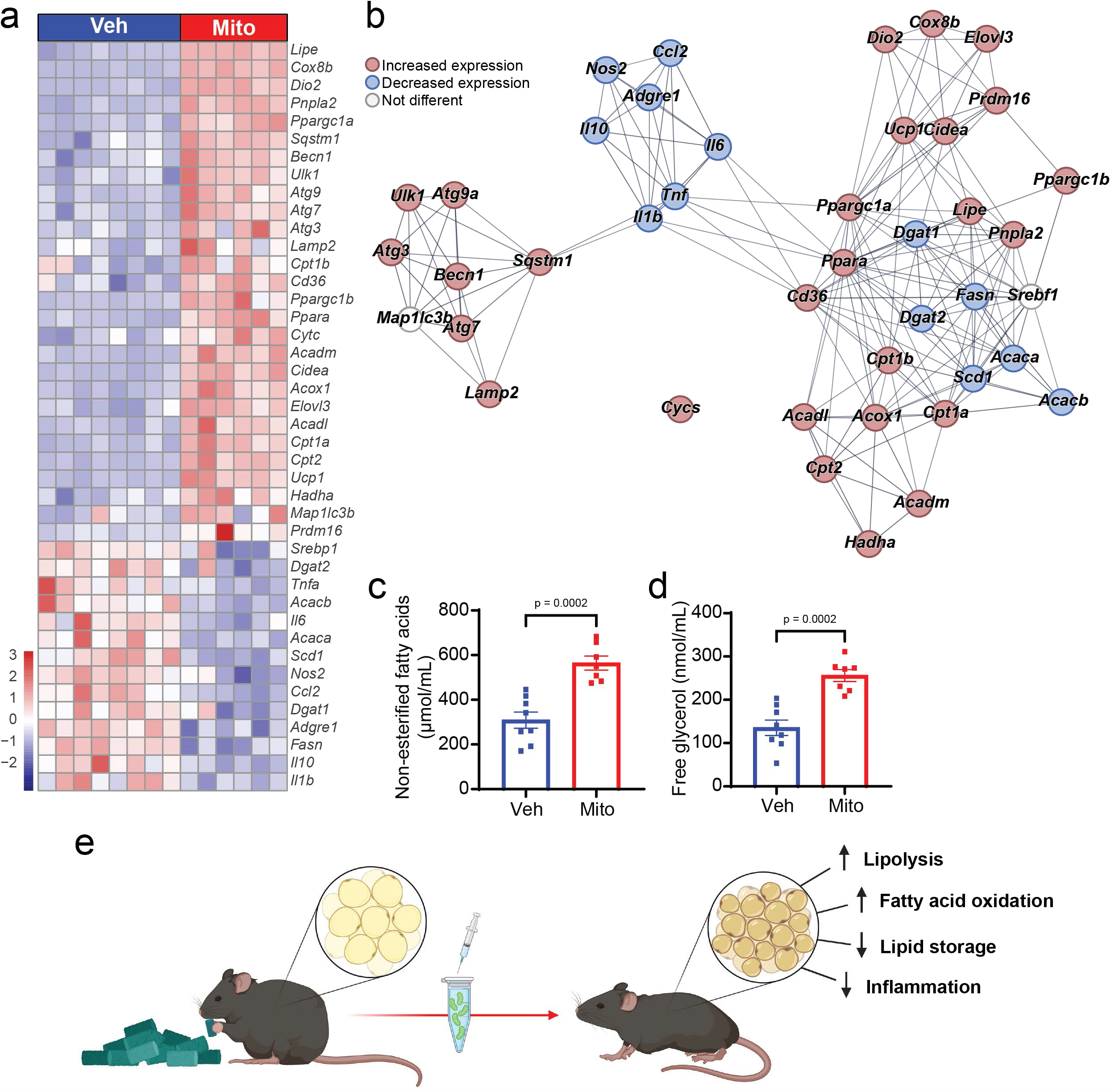
Transfer of exogenous mitochondria induces white adipose tissue adaptations that support systemic lipid oxidation and energy dissipation. Effect of mitochondrial transfer (Mito; red bars/points) compared to vehicle treatment (Veh; blue bars/points) on the expression of select transcripts in inguinal white adipose tissue (a) and their interaction map (b), and serum concentrations of non-esterified fatty acids (c) and free glycerol (d) as surrogate markers of lipolysis. Summary of observed relationship between mitochondrial transfer and adipose tissue adaptations (e). Values shown are the z-score (heatmap) and mean and standard error with biologically independent replicates overlayed (c-d; n=8 Veh and n=7 Mito). String clustering was performed with a pre-set interaction score of 0.7 (high confidence). Refer to each figure panel and Table 1 in the Extended Data for individual p-values. Source data are provided in the source data file. The image in panel e was created using BioRender.

All together, these data demonstrate that systemic transfer of exogenous mitochondria results in increased energy expenditure and enhanced lipid oxidation, in part, by inducing adaptations in white adipose tissue. We report that the magnitude of increased energy expenditure that results from mitochondrial transfer is capable of attenuating adiposity gains in mice with diet-induced obesity. Furthermore, we show that transfer of exogenous mitochondria restores systemic glucose tolerance and alleviates the accumulation of lipid in non-adipose tissues. This work is proof of principle that systemic administration of exogenous mitochondria is a viable strategy for reducing the deleterious effects of an obesogenic diet in mice.

## Discussion

In the past two decades, the discovery of adipose tissue plasticity in mice and humans has driven the search for potential druggable gene targets and pathways that reverse obesity and its comorbidities. Different genetic manipulations, as well as pharmacological, and environmental interventions have been suggested to improve mitochondrial function and promote the ‘browning’ or ‘beiging’ of white adipose tissue to facilitate an increase in energy expenditure^15,16^ Browning is marked by the presence of multilocular lipids in adipocytes, increased expression of transcripts involved in mitochondrial biogenesis and respiration, and a switch of white adipose tissue function from energy storage to energy expenditure, via increased uncoupling of respiration from ATP production^15^. However, chronic use of so-called browning agents has proven unfeasible due to their severe adverse effects, including hyperthermia, bone fractures and tachycardia^17^. Unlike previous attempts to increase energy expenditure through pharmacologic means, in the present study we were able to demonstrate that administration of exogenous mitochondria can cause changes in adipose tissue that promote systemic increases in energy expenditure, resulting in multiple metabolic benefits in the absence of any observable adverse effects.

The notion that the mitochondrial content of peripheral tissues is associated with overall metabolic health has been clearly demonstrated in both animal models and humans^18,19^. Furthermore, metabolic benefits of increasing mitochondrial content in adipose, muscle and other tissues have long been reported, primarily through exercise paradigms known to encourage mitochondrial biogenesis^20-25^. This association between mitochondrial volume and health outcomes suggests any treatment modality capable of increasing organ/tissue mitochondrial content is likely to have metabolic benefits that improve overall health. Here, we directly demonstrate that by increasing mitochondrial content via exogenous supply, adiposity gains, ectopic lipid deposition and impairments in glucose metabolism associated with diet-induced obesity can be mitigated. We also show that these metabolic improvements are due, in part, to the greater capacity exogenous mitochondria provide for supporting peripheral lipid oxidation and other energy consuming processes. These findings are important as they indicate that conditions of chronic bioenergetic stress can be overcome by providing additional cellular machinery to reduce the metabolic burden on existing systems.

The effects of treatment with exogenous mitochondria on energy expenditure in this study were, in part, due to an increase in total lipid oxidation supported by adaptations in white adipose tissue. However, we acknowledge the limitation of systemic administration on being able to clearly parse out tissue specific contributions. Since uptake of mitochondria by other tissues was also observed, we are unable to exclude the effects of administering exogenous mitochondria in other organ systems. Lungs, liver, kidney, brain, and skeletal muscle all take up exogenous mitochondria after systemic administration^7-14,26,27^ and all contribute to the regulation of energy balance. Indeed, given the large effect of mitochondrial transfer on lipid oxidation and the observed increase in dark photoperiod ambulation coincident with an increase in carbohydrate oxidation, it is hard to rule out the possibility that skeletal muscle, the main driver of lipid oxidation at rest and glucose use during exercise^28^, as a key tissue contributing to the energy balance phenotype we observed in this study. Future studies will assess direct routes of delivery to and quantitative assessment of mitochondrial uptake into these other tissues and allow us to determine the relevant contributions of each organ to the resulting improvements in metabolic flexibility.

In conclusion, we present proof of principle that transfer of exogenous mitochondria is an effective means of protecting against the deleterious effects of diet-induced obesity. Administering supplemental mitochondria can provide the additional bioenergetic support needed to increase energy expenditure, enhance oxidation of fatty acids, and sustain glucose tolerance and insulin sensitivity in the setting of diet-induced obesity. We also show that increased energy expenditure and enhanced lipid metabolism are mediated, in part, by transcriptional changes occurring in the white adipose tissue. Future studies will parse out the involvement of other tissues in driving these adaptations and will assess whether mitochondrial transfer can augment the effects of more traditional weight loss strategies.

## Methods

### Mitochondria Isolation, transfer, and Imaging

For *in vitro* experiments, mitochondria were isolated from HEK293 cells expressing a GFP targeted to mitochondria tag (addgene, plasmid #50057, Watertown, MA), whereas for *in vivo* experiments mitochondria were isolated from liver of *PhAM*^*excised*^ (Jackson Laboratory Stock No: 018397) or *PhAMfloxed* (Jackson Laboratory Stock No: 018385) crossed with *Alb1-cre* (Jackson Laboratory Stock No: 016832) mice using a previously described protocol^13^. Cells (or liver) were digested in ice-cold isolation buffer containing 300 nM sucrose, 10 mM HEPES, 1 mM EGTA and 2 mg/mL subtilisin A protease (Sigma-Aldrich, St. Louis, MO) and the homogenate was centrifuged at 500 x g for 5 min at 4°C. To separate mitochondria from other cellular debris, the supernatant was centrifuged at 800 x g for 10 min and again at 3000 x g for 10 min at 4°C. The mitochondrial pellet was reconstituted in cold sterile 1X PBS, and the protein concentration equivalent was quantified by Bradford assay. Functional viability of isolated mitochondria was assessed using CellTiter-Glo (Promega) to measure ATP concentrations. Mitochondrial uptake into organs was determined *ex vivo* using epifluorescent bioimaging (Xenogen IVIS Lumina, Perkin Elmer Akron, OH, USA) at 30 min, 2 h, 24 h, 48 h, or 72 h following a single 20 μg/g mitochondrial delivery.

### Cell culture

NIH**-**3T3-L1 cells (ATCC, Manassa, VA) were cultured in Dulbecco’s modified Eagle’s medium (DMEM) supplemented with 10% normal calf serum and 1% penicillin/streptomycin at 37°C, 5% CO_2_ until confluency. On Day 0, cells were induced to differentiate into mature adipocytes in DMEM containing 10% Fetal bovine serum (FBS) and a cocktail of 0.5 μM isobutyl-1-methylxanthine, 1 μM dexamethasone and 1.7 μM insulin. After 48 h, cells were changed to 10% FBS/DMEM and insulin, which was refreshed every two days until cells were fully differentiated on Day 8. Under all experimental conditions, cells were treated with either 10 μg of mitochondria or a matched volume of 1x PBS.

### Quantification of mitochondrial DNA

Total DNA was extracted from NIH-3T3-L1 cells by incubating cells in SDS lysis buffer with proteinase K overnight at 55°C followed by ethanol-precipitation. The abundance of mitochondrial DNA (mtDNA) was quantified using real-time polymerase chain reaction (RT-PCR) by assessing the ratio of human MT-NADH dehydrogenase, subunit 6 (ND6) relative to mouse MT-NADH dehydrogenase, subunit 1 (ND1). Primer sequences are provided in the Extended Data Table 2.

### Cellular respiration

Oxygen consumption rate (OCR) in NIH-3T3-L1 adipocytes treated with exogenous mitochondria was measured using the Seahorse XF24 Extracellular Flux Analyzer (Agilent, Santa Clara, CA, USA) with slight modifications of the manufacturer’s protocol. Briefly, the adipocytes were incubated in DMEM with an adjusted pH value of 7.4 at 37^?^C for 1 h. After measuring basal OCR, mitochondrial stress tests were performed by adding all reagents to the adipocytes through the ports sequentially as follows: oligomycin (2 μM), carbonyl cyanide 4-trifluoromethoxy, phenylhydrazone (FCCP; 2 μM), and antimycin A (0.5 μM)/ rotenone (0.5 μM).

### Lipolysis

NIH-3T3-L1 cells were incubated in DMEM containing 2% fatty-acid free BSA in the presence of exogenous mitochondria and/or CL316,243 (β_3_-adrenergic receptor agonist; Sigma Aldrich) for 72 h. Media was collected from cultured cells and free glycerol (Sigma Aldrich) and non-esterified fatty acids (NEFA) (ZenBio) were determined using kits according to manufacturer’s protocols.

### Animal studies

All animal procedures were approved by the University of Tennessee Health Science Center Institutional Animal Care and Use Committee. Male C57BL/6J mice were purchased from Jackson Laboratories. Mice were housed on a 12-h light/12-h dark cycle with free access to food and water. Beginning at 8 weeks of age, mice were provided a high fat diet to promote obesity (60% Kcal fat, Research Diets D12492, New Brunswick, NJ). From 14 weeks of age (after 6 weeks of high fat diet) mice began receiving weekly retro-orbital injections containing either mitochondria isolated from a lean donor mouse (20 μg/g body weight) or a matched volume of vehicle (sterile PBS).

### Glucose tolerance, insulin responsiveness, and lipolysis

After 12 weeks of treatment with either mitochondria or vehicle, mice were fasted for 6 h, and blood glucose was measured using a handheld glucometer (Contour Next EZ, Parsipanny, NJ). For glucose tolerance testing, mice were injected with glucose intraperitoneally (2 g/kg body weight) and glucose was measured from tail blood at regular intervals over a 2 h period. To determine insulin responsiveness, mice were given an intraperitoneal injection of insulin (1 U/Kg body weight), and blood glucose was measured every 15 min over one hour. As surrogate markers for rates of basal lipolysis, the concentrations of free glycerol and non-esterified fatty acids were determined in serum obtained from mice after 15 weeks of treatment as described above.

### Body composition, indirect calorimetry, activity, and energy intake

Mice were weighed weekly and body composition was measured every four weeks by magnetic resonance (EchoMRI, Echo Medical Systems, Houston, TX). After 13 weeks of treatment, mice were individually housed in a high-resolution home cage-style Comprehensive Laboratory Animal Monitoring System capable of measuring respiratory gases via open-circuit indirect calorimetry, physical activity by infrared beam breaks in the x- and y-axes, and food intake via load cell-linked hanging feed baskets (Columbus Instruments, Columbus, OH). Energy expenditure^29^ and carbohydrate and lipid oxidation^30,31^ were calculated from VO_2_ and VCO_2_ values (Columbus Instruments, Columbus, OH). Total physical activity was calculated as the combined number of beam breaks along the x- and y-axes, whereas ambulatory activity was calculated as the combined number of consecutive x- and y-axes beam breaks occurring in a single series. The difference in beam breaks between ambulatory and total activity was considered stereotypic activity (i.e., rearing, grooming, etc.). Resting energy expenditure was calculated from data acquired during ZT 0-4 during periods where mice were not feeding and not moving. The difference between total and resting energy expenditure was considered non-resting energy expenditure. Energy intake was determined from individual feeding bouts after which cumulative daily food intake was determined and energy intake calculated using the density of metabolizable energy in the diet as provided by the manufacturer. Data for each mouse was collected over 7 days under ambient housing conditions (22°C). Data from the first day was discarded to account for acclimation to the change in housing and for every subsequent day, data were binned by hour and the hourly mean values (indirect calorimetry data) or summed values (activity and food intake data) over each 24-h measurement period were used to determine the reported mean hourly values for each mouse. These data were analyzed using mixed linear models with total body weight, body fat and lean mass included as random effect variables (time course data), and by ANCOVA with fat mass and lean mass included as covariates (daily values), as previously described^32^. Estimated marginal means were calculated with both fat and lean mass as constants in the model. All figures except that of the estimated marginal means show absolute (non-mass adjusted) individual and mean data ± standard error.

### Histology

Cells were fixed in 10% neutral buffered formalin for 20 min followed by either Oil Red O and hematoxylin or BODIPY493/503 and DAPI for 20 min at room temperature to assess lipid content. Liver and adipose tissue were isolated from mice after 15 weeks of treatment, fixed in 10% neutral buffered formalin, and dehydrated in ethanol. Samples were embedded in paraffin and 5 μm sections were cut and stained with hematoxylin and eosin (H&E) All images were captured using the EVOS FLoid imaging system microscope (Thermo Fisher Scientific). Adipocyte size was determined using Image J (https://imagej.nih.gov/ij/).

### RNA isolation and q-RT-PCR

Cells and tissues were collected in Trizol (Thermo Fisher Scientific), and total RNA isolated following the manufacturer’s protocol. RNA was reverse transcribed using the using the High-Capacity cDNA Kit (Applied Biosystems). Quantitative RT-PCR was performed using PowerUp SYBR green master mix (Applied Biosystems) on a QuantStudio 6 Flex system. Relative expression of each transcript of interest was calculated using the ΔΔCt method with *CypA* as an internal housekeeping reference. Primer sequences are provided in the Extended Data Table 2.

### Statistical analyses

Cell culture experiments were independently replicated a minimum of 3 times and all treatment conditions performed in triplicate. Animal studies included a minimum of 7 mice per group. Data were analyzed and visualized using either GraphPad Prism 9 (GraphPad Software, La Jolla California USA) or R version 4.1.2 and the following packages: tidyverse 1.3.1, lubridate 1.8.0, data.table 1.14.2, lme4 1.1-28, emmeans 1.7.2, rstatix 0.7.0, car 3.0-12, and pheatmap 1.0.12. Statistical significance (p- or q-values) was set *a priori* to <0.05. All data are presented as mean ± standard error of the mean unless otherwise specified. Unpaired 2-tailed Student’s *t* tests, Welch’s *t* tests, Wilcoxon-Rank-Sum tests, one-way analysis of variance (ANOVA) followed by posthoc Tukey’s multi-comparison tests, Analysis of Co-variance (ANCOVA), or mixed linear models with likelihood ratio tests were used as appropriate. For specific details about statistical tests used, refer to the figure legends.

## Acknowledgments

This work was supported by funds from the National Institutes of Health grants R01DK117183 (AB) and R01DK132230 (NSP and AB), This study was presented, in part, as an oral abstract at Obesity Week 2023 (ES). Schematic diagrams were generated using BioRender.

## Author contributions

MN, AB, AS, and EJS designed experiments. SM, AS, CW, CW, and PSP performed experiments. MN, AB, and EJS analyzed and interpreted data. MN, AS, AB, and EJS prepared figures. MN, AB, and EJS wrote the manuscript. AB, EJS, MN, PSP, KM, and NP edited and/or provided feedback on the manuscript. All authors approved submission of the manuscript.

