## Extended Data for "Exogenous mitochondrial transfer increases energy expenditure and attenuates adiposity gains in mice with diet-induced obesity"

***Extended Data Figure 1: Transfer of exogenous mitochondria increases respiration, enhances lipolysis, and reduces lipogenesis in vitro***

Mitochondrial dose response curve in NIH-3T3-L1 cells after 24 h of exposure to exogenous mitochondria (a). $O_2$ consumption rate (OCR) of NIH-3T3-L1 cells during a mitochondrial stress test (b) and quantitation of basal OCR (c). Schematic depicting differentiation of NIH-3T3-L1 cells into adipocytes (d). Representative images showing lipid staining of NIH-3T3-L1 adipocytes following 72 h incubation with either vehicle (Veh) or 10 µg/mL HEK293 mitochondria (Mito; e). Heatmap depicting the effect of mitochondrial incubation on the expression of transcripts associated with respiration, lipolysis, fatty acid oxidation, lipogenesis, adipogenesis, and inflammation in NIH-3T3-L1 adipocytes after 72 h (f). Non-esterified fatty acids (g) and free glycerol concentrations (h) in media under basal and CL316243 stimulated conditions. Values shown are the mean and standard error with independent replicates overlayed (a-c, g) or z-scores (f). Each experiment depicts the results of three independent experiments (n=3 Veh and n=3 Mito). Refer to each figure panel for individual p-values. The image in panel d was created using BioRender. Abbreviations: OCR, $O_2$ consumption rate; DAPI, 4',6-diamidino-2-phenylindole; BODIPY, 4,4-difluoro-4-bora-3a,4a-diaza-s-indacene; Veh, vehicle, Mito, mitochondria, Stim., Stimulated with CL316243.

***Extended Data Figure 2: Systemic transfer leads to uptake of exogenous mitochondria into multiple organs***

Epifluorescence of a selection of organs from mice injected with 20 µg/g body weight fluorescently labelled exogenous mitochondria 30 min, 2 h, 24 h, 48 h, and 72 h post-injection. Schematic depicting experimental design (a). Brain (b), triceps surae (c), quadriceps (d), liver (e), kidneys (f), interscapular BAT (g), heart (h), and lung (i). The image in panel a was created with BioRender.com.

### Extended Data Figure 1

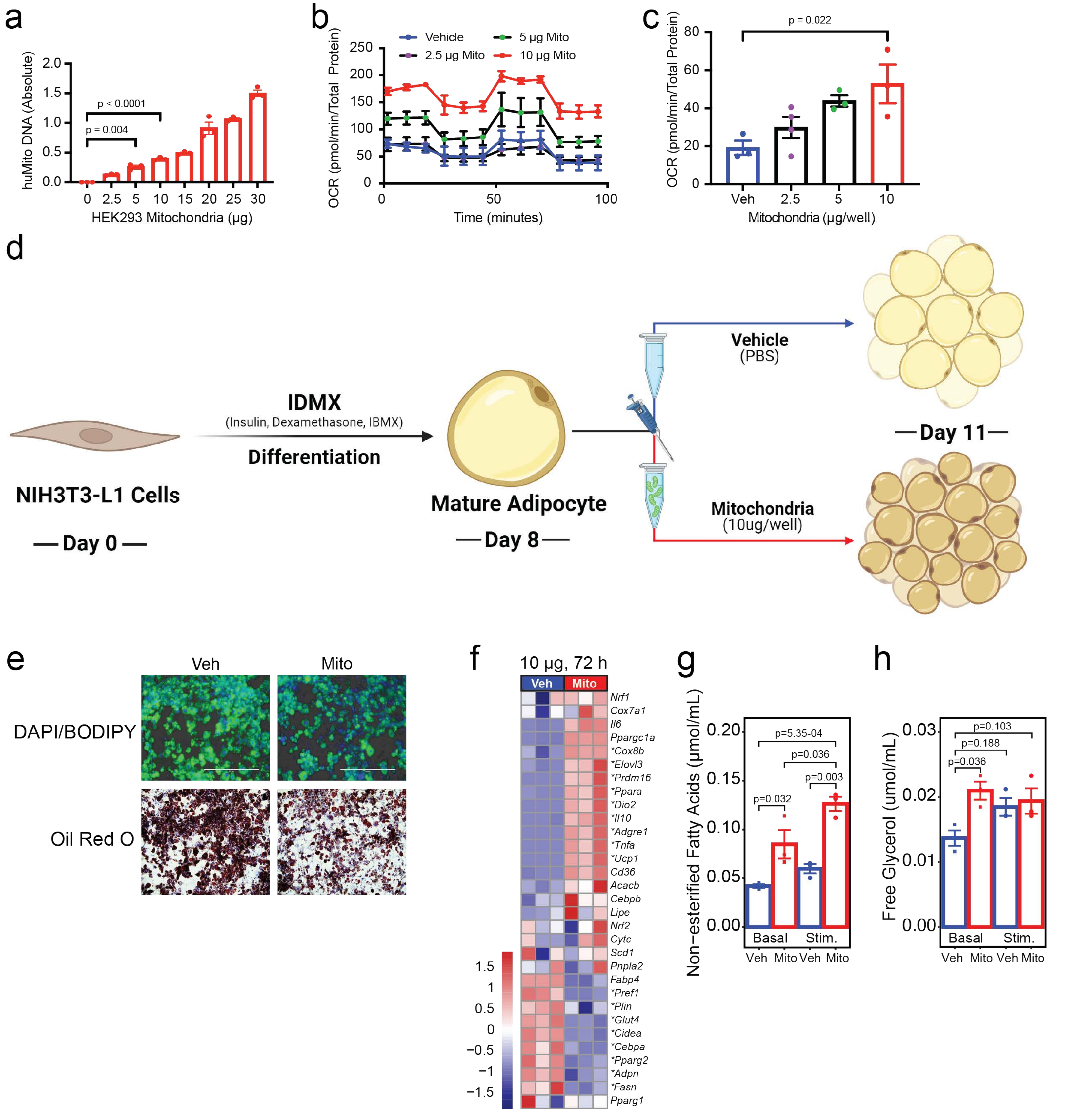

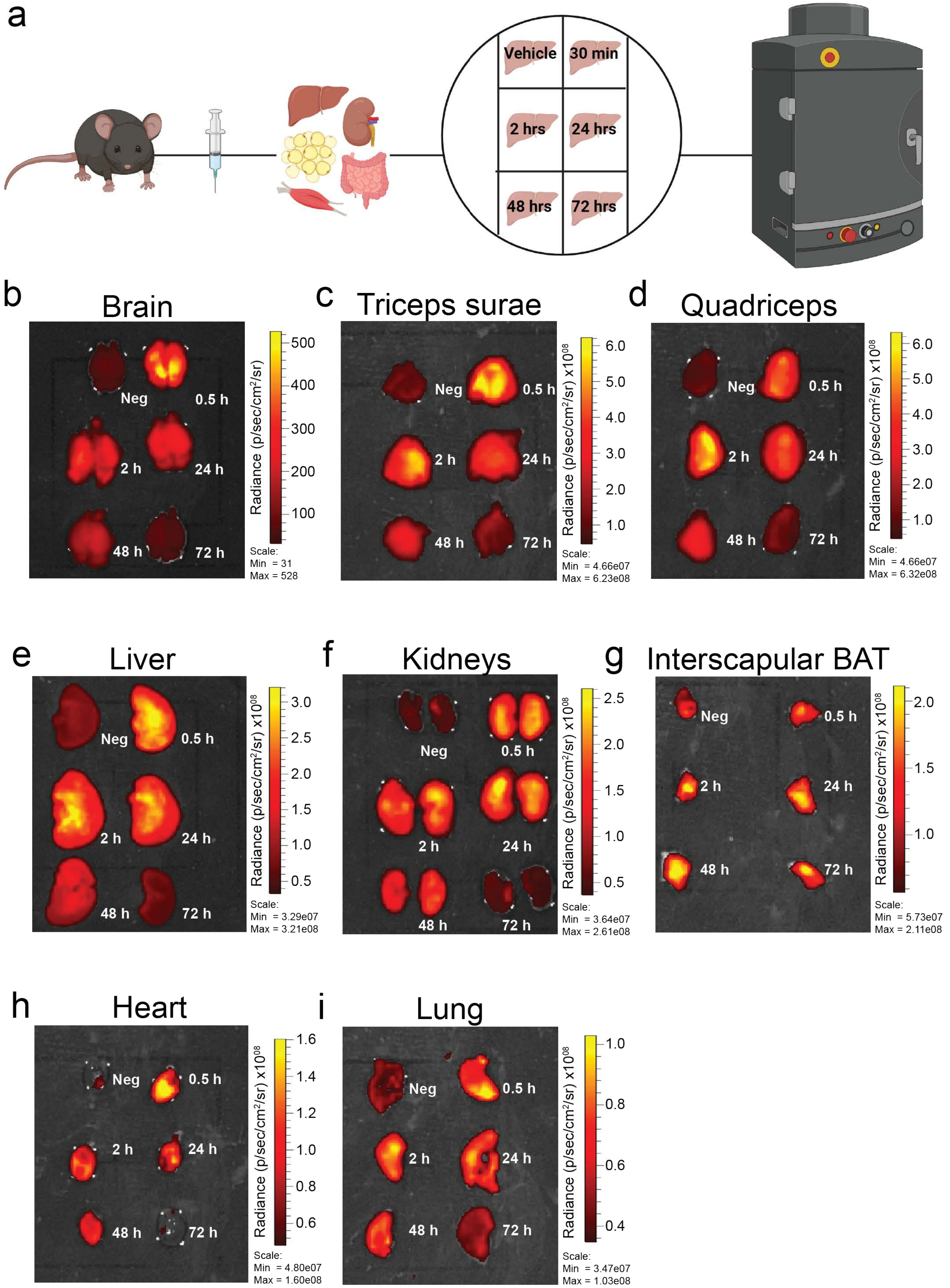
a
Vehicle
30 min
2 hrs
24 hrs
48 hrs
72 hrs
b
Brain
Neg
0.5 h
2 h
24 h
48 h
72 h
Radiance (p/sec/cm²/sr)
500
400
300
200
100
Scale:
Min = 31
Max = 528
c
Triceps surae
Neg
0.5 h
2 h
24 h
48 h
72 h
Radiance (p/sec/cm²/sr) x10⁰⁸
6.0
5.0
4.0
3.0
2.0
1.0
Scale:
Min = 4.66e07
Max = 6.23e08
d
Quadriceps
Neg
0.5 h
2 h
24 h
48 h
72 h
Radiance (p/sec/cm²/sr) x10⁰⁸
6.0
5.0
4.0
3.0
2.0
1.0
Scale:
Min = 4.66e07
Max = 6.32e08
e
Liver
Neg
0.5 h
2 h
24 h
48 h
72 h
Radiance (p/sec/cm²/sr) x10⁰⁸
3.0
2.5
2.0
1.5
1.0
0.5
Scale:
Min = 3.29e07
Max = 3.21e08
f
Kidneys
Neg
0.5 h
2 h
24 h
48 h
72 h
Radiance (p/sec/cm²/sr) x10⁰⁸
2.5
2.0
1.5
1.0
0.5
Scale:
Min = 3.64e07
Max = 2.61e08
g
Interscapular BAT
Neg
0.5 h
2 h
24 h
48 h
72 h
Radiance (p/sec/cm²/sr) x10⁰⁸
2.0
1.5
1.0
Scale:
Min = 5.73e07
Max = 2.11e08
h
Heart
Neg
0.5 h
2 h
24 h
48 h
72 h
Radiance (p/sec/cm²/sr) x10⁰⁸
1.6
1.4
1.2
1.0
0.8
0.6
Scale:
Min = 4.80e07
Max = 1.60e08
i
Lung
Neg
0.5 h
2 h
24 h
48 h
72 h
Radiance (p/sec/cm²/sr) x10⁰⁸
1.0
0.8
0.6
0.4
Scale:
Min = 3.47e07
Max = 1.03e08

Table 1: qPCR results

| | Inguinal white adipose depot | | | | Epididymal white adipose depot | | | | NIH-3T3-L1 adipocytes | | | | |
|---|---|---|---|---|---|---|---|---|---|---|---|---|---|
| **Transcript** | **Veh (T)** | **Mito (T)** | **p value** | **q value** | **Veh (T)** | **Mito (T)** | **p value** | **q value** | **Transcript** | **Veh (72 hrs)** | **Mito (72 hrs** | **P value** | **q value** |
| *Ppara* | 1.001 | 1.613 | 0.000001 | 0.000001 | 1.008 | 2.265 | 0.000138 | 0.000056 | *Ppara* | 1.009 | 8.853 | 0.003161 | 0.00329 |
| *Ucp1* | 1.005 | 3.522 | <0.000001 | <0.000001 | 1.016 | 4.028 | 0.008516 | 0.001966 | *Ucp1* | 1.011 | 30.53 | 0.001069 | 0.00174 |
| *Dio2* | 1.003 | 2.627 | <0.000001 | <0.000001 | 1.032 | 1.015 | 0.901738 | 0.173477 | *Dio2* | 1.019 | 15.66 | 0.003258 | 0.00329 |
| *Ppargc1a* | 1.009 | 2.267 | <0.000001 | <0.000001 | 1.015 | 2.518 | <0.000001 | <0.000001 | *Ppargc1a* | 1.003 | 8.913 | <0.000001 | <0.000001 |
| *Ppargc1b* | 1.01 | 3.402 | 0.000096 | 0.000096 | 1.007 | 3.352 | 0.000607 | 0.000182 | *Prdm16* | 1.012 | 33.54 | 0.005965 | 0.005086 |
| *Prdm16* | 1.015 | 2.595 | 0.019809 | 0.019809 | 1.063 | 2.882 | 0.003075 | 0.000776 | *Elovl3* | 1.002 | 2.157 | 0.00792 | 0.006315 |
| *Elovl3* | 1.106 | 3.051 | 0.000012 | 0.000012 | 1.012 | 3 | 0.000049 | 0.000023 | *Cidea* | 1.001 | 0.5557 | 0.000286 | 0.001318 |
| *Cidea* | 1.006 | 3.279 | <0.000001 | <0.000001 | 1.011 | 1.408 | 0.000371 | 0.000124 | *Cox8b* | 1.011 | 1.922 | 0.001053 | 0.00174 |
| *Cox8b* | 1.001 | 2.887 | <0.000001 | <0.000001 | 1.001 | 2.06 | <0.000001 | <0.000001 | *Cycs* | 1.003 | 1.112 | 0.469299 | 0.263329 |
| *Cycs* | 1.002 | 1.232 | 0.000133 | 0.000133 | 1.005 | 1.524 | 0.00001 | 0.000005 | *Lipe* | 0.9999 | 1.106 | 0.151479 | 0.095621 |
| *Lipe* | 1.006 | 2.025 | <0.000001 | <0.000001 | 1.013 | 2.097 | 0.000005 | 0.000003 | *Pnpla2* | 1.001 | 0.9843 | 0.800849 | 0.433316 |
| *Pnpla2* | 1.006 | 2.392 | <0.000001 | <0.000001 | 1.013 | 2.775 | <0.000001 | <0.000001 | *Fasn* | 1.021 | 0.4312 | 0.019517 | 0.014784 |
| *Fasn* | 1.003 | 0.6597 | 0.000059 | 0.000059 | 1.017 | 0.6492 | 0.000589 | 0.000182 | *Scd1* | 1.057 | 1.037 | 0.940115 | 0.459443 |
| *Scd1* | 1.008 | 0.6894 | 0.000216 | 0.000216 | 1.007 | 0.5517 | <0.000001 | <0.000001 | *Cd36* | 1.002 | 6.931 | 0.000448 | 0.001358 |
| *Cd36* | 1.006 | 1.294 | 0.000253 | 0.000253 | 1.007 | 2.296 | <0.000001 | <0.000001 | *Acacb* | 1.001 | 1.398 | 0.053662 | 0.036954 |
| *Dgat1* | 1.003 | 0.8328 | 0.012065 | 0.012065 | 1.009 | 0.8205 | 0.01127 | 0.002465 | *Il6* | 1.017 | 14.33 | 0.000538 | 0.001358 |
| *Dgat2* | 1.002 | 0.8232 | 0.007727 | 0.007727 | 1.009 | 2.258 | <0.000001 | <0.000001 | *Tnfa* | 1.004 | 22.45 | 0.001638 | 0.002068 |
| *Acaca* | 1.026 | 0.5701 | 0.001692 | 0.001692 | 1.008 | 0.7195 | 0.000191 | 0.000067 | *Il10* | 1.016 | 12.05 | 0.00117 | 0.00174 |
| *Acacb* | 1.034 | 0.4989 | 0.001309 | 0.001309 | 1.022 | 0.8084 | 0.082174 | 0.017025 | *Adgre1* | 1.01 | 35.36 | 0.001198 | 0.00174 |
| *Srebp1* | 1.003 | 0.7899 | 0.024248 | 0.024248 | 1.005 | 0.8979 | 0.104656 | 0.02114 | *Adipoq* | 1.001 | 0.6602 | 0.002618 | 0.003051 |
| *Cpt2* | 1.001 | 1.924 | <0.000001 | <0.000001 | 1.004 | 1.971 | <0.000001 | <0.000001 | *Pparg1* | 1.011 | 0.9948 | 0.888155 | 0.459443 |
| *Cpt1a* | 1.006 | 2.428 | <0.000001 | <0.000001 | 1.003 | 1.284 | 0.000024 | 0.000012 | *Pparg2* | 1.008 | 0.4349 | 0.004003 | 0.00379 |
| *Cpt1b* | 1.031 | 1.523 | 0.008707 | 0.008707 | 1.025 | 2.945 | 0.000001 | <0.000001 | *Cebpa* | 1.013 | 0.3826 | 0.006043 | 0.005086 |
| *Acox1* | 1.002 | 1.742 | 0.000002 | 0.000002 | 1.005 | 1.765 | 0.000062 | 0.000026 | *Cebpb* | 1 | 1.161 | 0.082231 | 0.054165 |
| *Hadha* | 1.024 | 1.801 | 0.000276 | 0.000276 | 1.014 | 1.788 | 0.000003 | 0.000002 | *Pref1* | 1.004 | 0.4831 | 0.001264 | 0.00174 |
| *Acadl* | 1.004 | 1.746 | 0.000013 | 0.000013 | 1.009 | 1.666 | 0.000009 | 0.000005 | *Fabp4* | 1 | 0.5668 | 0.000348 | 0.001318 |
| *Acadm* | 1.001 | 1.956 | <0.000001 | <0.000001 | 1.015 | 1.658 | 0.000061 | 0.000026 | *Glut4* | 1.001 | 0.6675 | 0.0003 | 0.001318 |
| *Ccl2* | 1.005 | 0.7778 | 0.000342 | 0.000342 | 1.01 | 0.7546 | 0.002722 | 0.00071 | *Plin* | 1.001 | 0.7159 | 0.031121 | 0.022451 |
| *Il6* | 1.036 | 0.5211 | 0.002894 | 0.002894 | 1.012 | 0.6911 | 0.001433 | 0.000414 | *Cox7a1* | 1.001 | 1.075 | 0.214891 | 0.130224 |
| *Tnfa* | 1.022 | 0.6985 | 0.007523 | 0.007523 | 1.014 | 0.7499 | 0.001971 | 0.000549 | *Nrf1* | 1.005 | 1.102 | 0.256279 | 0.149332 |
| *Il10* | 1.022 | 0.6127 | 0.001806 | 0.001806 | 1.017 | 0.7138 | 0.011289 | 0.002465 | *Nrf2* | 1.002 | 1.013 | 0.938839 | 0.459443 |
| *Adgre1* | 1.001 | 0.786 | 0.00005 | 0.00005 | 1.016 | 0.7981 | 0.012217 | 0.002598 | | | | | |
| *Nos2* | 1.002 | 0.7752 | 0.001512 | 0.001512 | 1.002 | 0.8129 | 0.000183 | 0.000067 | | | | | |
| *Il1b* | 1.004 | 0.8632 | 0.005394 | 0.005394 | 1.018 | 0.6381 | 0.002198 | 0.000592 | | | | | |
| *Atg3* | 1.001 | 1.334 | 0.000847 | 0.000847 | 1.006 | 1.412 | <0.000001 | <0.000001 | | | | | |
| *Sqstm1* | 1.011 | 1.58 | 0.000007 | 0.000007 | 1.002 | 1.289 | 0.00016 | 0.000062 | | | | | |
| *Ulk1* | 1.02 | 1.799 | 0.000035 | 0.000035 | 1.008 | 1.354 | 0.005107 | 0.001214 | | | | | |
| *Becn1* | 1.012 | 1.651 | 0.000015 | 0.000015 | 1.007 | 1.568 | 0.000001 | <0.000001 | | | | | |
| *Map1lc3b* | 1.008 | 1.259 | 0.024023 | 0.024023 | 1.01 | 1.402 | 0.000382 | 0.000124 | | | | | |
| *Atg7* | 1.008 | 1.64 | 0.000018 | 0.000018 | 1.01 | 1.626 | <0.000001 | <0.000001 | | | | | |
| *Atg9* | 1.002 | 1.57 | 0.000011 | 0.000011 | 1.008 | 1.637 | 0.003646 | 0.000893 | | | | | |
| *Lamp2* | 1.001 | 1.202 | 0.008498 | 0.008498 | 1.005 | 0.9581 | 0.422765 | 0.083316 | | | | | |

Table 2: Primer sequences

| Transcript/Gene | Forward (5'-->3') | Reverse (5'-->3") |
|---|---|---|
| *Il10* | CAG AGC CAC ATG CTC CTA GA | GTC CAG CTG GTC CTT TGT TT |
| *Nos2* | CGA AAC GCT TCA CTT CCA A | TGA GCC TAT ATT GCT GTG GCT |
| *Il1b* | AAT GAC CTG TTC TTT GAA GTT GAC | GTG ATA CTG CCT GCC TGA AG |
| *Tnfa* | CCT CCC TCT CAT CAG TTC TAT GG | CGT GGG CTA CAG GCT TGT C |
| *Il6* | TGG CTA AGG ACC AAG ACC ATC CAA | AAC GCA CTA GGT TTG CCG AGT AGA |
| *Ccl2* | TCA CCT GCT GCT ACT CAT TCA CCA | TAC AGC TTC TTT GGG ACA CCT GCT |
| *Becn1* | ATG GAG GGG TCT AAG GCG TC | TCC TCT CCT GAG TTA GCC TCT |
| *Atg7* | GTT CGC CCC CTT TAA TAG TGC | TGA ACT CCA ACG TCA AGC GG |
| *Atg9* | CAG TTT GAC ACT GAA TAC CAG CG | AAT GTG GTG CCA AGG TGA TTT |
| *Lamp2* | TGT ATT TGG CTA ATG GCT CAG C | TAT GGG CAC AAG GAA GTT GTC |
| *Srebf1* | GAT GTG CGA ACT GGA CAC AG | CAT AGG GGG CGT CAA ACA G |
| *Scd1* | GAG TCA CAA GAG TAG CTG AG | ATC ATT AAC ACC CCG ATA GC |
| *Cd36* | GTC TGA AGG ACT GGA ATG C | GGG TCT CAA CCA TTC ATC TAT |
| *Hadha* | TGA AAA CAA GCA ATG TGG CTA | TGA AGA GAT ACA AGC CAT GGT G |
| *Cidea* | TGC TCT TCT GTA TCG CCC AGT | GCC GTG TTA AGG AAT CTG CTG |
| *Dio2* | AGA GTG GAG GCG CAT GCT | GGC ATC TAG GAG GAA GCT GTT C |
| *Elovl3* | CCA ACA ACG ATG AGC AAC AG | CGG GTT AAA AAT GGA CCT GA |
| *Ppara* | GGG TAC CAC TAC GGA GTT CAC G | CAG ACA GGC ACT TGT GAA AAC G |
| *Ucp1* | ACT GCC ACA CCT CCA GTC ATT | CTT TGC CTC ACT CAG GAT TGG |
| *Fasn* | GGA GGT GGT GAT AGC CGG TAT | TGG GTA ATC CAT AGA GCC CAG |
| *Acaca* | ATG GGC GGA ATG GTC TCT TTC | TGG GGA CCT TGT CTT CAT CAT |
| *Acacb* | CGC TCA CCA ACA GTA AGG TGG | GCT TGG CAG GGA GTT CCT C |
| *Dgat1* | TCC GTC CAG GGT GGT AGT G | TGA ACA AAG AAT CTT GCA GAC GA |
| *Dgat2* | GCG CTA CTT CCG AGA CTA CTT | GGG CCT TAT GCC AGG AAA CT |
| *Pnpla2* | AAC ACC AGC ATC CAG TTC AA | GGT TCA GTA GGC CAT TCC TC |
| *Cox8b* | GAA CCA TGA AGC CAA CGA CT | GCG AAG TTC ACA GTG GTT CC |
| *Cycs* | TCC ATC AGG GTA TCC TCT CC | GGA GGC AAG CAG AAG ACT GG |
| *Lipe* | CCT GCA AGA GTA TGT CAC GC | GGA GAG AGT CTG CAG GAA CG |
| *Ppargc1a* | TGA AAG AAG CGG TGA ACC ACT G | TGG CAT CTC TGT GTC AAC CAT G |
| *Ppargc1b* | GCA TGG TGC CTT CGC TGA | TGG CAT CTC TGT GTC AAC CAT G |
| *Atg3* | ACA CGG TGA AGG GAA AGG C | TGG TGG ACT AAG TGA TCT CCA G |
| *Sqstm1* | AGG ATG GGG ACT TGG TTG C | TCA CAG ATC ACA TTG GGG TGC |
| *Ulk1* | AAG TTC GAG TTC TCT CGC AAG | CGA TGT TTT CGT GCT TTA GTT CC |
| *Adgre1* | TGA CTC ACC TTG TGG TCC TAA | CTT CCC AGA ATC CAG TCT TTC C |
| *Cpt1a* | TTT GAA TCG GCT CCT AAT GG | CCC AAG TAT CCA CAG GGT CA |
| *Prdm16* | CAG CAC GGT GAA GCC ATT C | GCG TGC ATC CGC TTG TG |
| *Acox1* | CTT CGA GGG GGA GAA CAC T | CCC GAC TGA ACC TGG TCA TA |
| *Acadl* | ATG GCT GCG CGC CTG CTC CTC | TGG ATT CTC AAT GGA AGC AAG |
| *Acadm* | GGA TGA CGG AGC AGC CAA TG | ATA CTC GTC ACC CTT CTT CT |
| *Cpt1b* | CCC GAG CAG TGC CGG GAA GC | GAA ATG AGC CAG CTG TAG GG |
| *human mt-ND6* | CCC CAC AAA CCC CAT TAC TAA ACC CA | TTT CAT CAT GCG GAG ATG TTG GAT GG |
| *mouse mt-ND1* | CTAGCAGAAACAAACCGGGC | CCGGCTGCGTATTCTACGTT |
